# An experimental test of the influence of microbial manipulation on sugar kelp (*Saccharina latissima*) supports the the core influences host function hypothesis

**DOI:** 10.1101/2025.02.26.640414

**Authors:** Jungsoo Park, Evan Kohn, Siobhan Schenk, Katherine Davis, Jennifer Clark, Laura Wegener Parfrey

## Abstract

Kelp are valued for a wide range of commercial products and for their role in kelp forest ecosystems, making kelp cultivation a rapidly expanding economic sector. Microbes associated with kelp and other macroalgae play a critical role in processes such as nutrient exchange, chemical signaling, and defense against pathogens. Thus, manipulating the microbiome to enhance macroalgal growth and resilience is a promising yet underexplored approach for sustainable kelp cultivation. The core microbiome hypothesis suggests that the bacteria that are consistently found on a host (the core microbes) are likely to have a disproportionate impact on host biology, making them an attractive target for microbiome manipulation. In this study, we surveyed wild *Saccharina latissima* and their surrounding environment to identify core bacterial taxa, compared them to cultivated kelp, and experimentally tested how cultured bacterial isolates affect kelp development. We found that core bacteria are nearly absent in cultivated juvenile sporophytes in nurseries but eventually colonized them after outplanting to ocean farm sites. Bacterial inoculants had both positive and negative effects on kelp development. Notably, the strength of association of a bacterial genus with kelp in the wild positively correlated with its impact on gametophyte settlement and sporophyte development in kelp co-culture experiments, aligning with predictions from the core microbiome influences host function hypothesis. These findings affirm the feasibility of using microbial manipulations to improve current kelp aquaculture practices and provide a framework for developing these techniques.

**IMPORTANCE:** Microorganisms consistently associated with hosts are widely thought to be more likely to impact host biology and health. However, this intuitive concept has not been experimentally evaluated. This study formalizes this concept as the Core microbiome Influences Host Function hypothesis and experimentally tests this hypothesis in sugar kelp (*Saccharina*). The distribution of bacteria on wild kelp and core microbes were first identified by compiling a broad dataset of the kelp microbiome sampled across space and time. Bacterial cultures were isolated from the surface of sugar kelp and individually grown in laboratory co-culture with sugar kelp spores to assess the ability of bacterial isolates to influence kelp growth and development. In support of the core influences host function hypothesis, isolates belonging to bacterial genera that are more strongly associated with wild sugar kelp are more likely to influence development in laboratory experiments.

## INTRODUCTION

The kelp *Saccharina latissima* (Phaeophyceae) is one of the most important species for commercial aquaculture in the northern hemisphere. *Saccharina* has been explored as a focal species for biofuels, livestock feed, and as a component of integrated multi-trophic aquaculture (1–3) due to its fast growth and its richness in valuable organic compounds (4–6). Globally, increasing consumer demand for new protein sources and climate friendly foods (7) has also been a key driver for the expansion of macroalgal cultivation outside of Asia (8,9). Expanding kelp cultivation in North America not only relies on finding a suitable market, but also requires innovation to reduce cost and increase reliability, especially in the land-based nursery phase (10). Moreover, kelp cultivation faces growing challenges associated with rising global temperatures (11,12) and increasing disease pressure (13). The intensification and global expansion of macroalgal cultivation exacerbate these challenges, as does climate change, leading to increased disease pressure that reduces crop yields (14–16). As such, developing strategies to mitigate these pressures is crucial for supporting the developing kelp aquaculture industry.

Recent studies have highlighted the importance of symbiotic bacteria for host development, survival, and fitness (17–20). This has prompted a wave of interest in manipulating the microbiome to benefit hosts, targeting taxa that may increase crop productivity or resistance to microbial pathogens (21,22). One microbiome manipulation strategy is the addition of microbes with beneficial effects, also called probiotics. In agriculture, addition of plant-growth promoting bacteria has been shown to reduce the need for chemical fertilizers (23). For example, *Rhizobia* spp. and several free-living bacteria, such as *Azospirillum* spp., have the capability to fix nitrogen which is then utilized by plants (24). *Pseudomonas putida* GR12-2 produces indole-3-acetic acid (IAA), the most commonly occuring form of the phytohormone, auxin, which influences various aspects of plant growth, including cell division, extension, and differentiation (25). Similarly, addition of specific bacterial taxa can increase growth rates of microalgae (e.g. *Tetraselmis* species) and macroalgae (e.g. *Ulva*) (26–28). The addition of bacterial inoculants in lab settings also influences tolerance to low salinity stress (29) and disease susceptibility (30) in *Ulva* sp., *Ectocarpus* sp., and *Gracilaria* sp. (31–34). Hence, manipulation of the microbiome in macroalgal aquaculture presents a promising but underexplored strategy (35).

Harnessing the microbiome for kelp aquaculture requires an understanding of the kelp cultivation process, and how microbes influence this process. Commercial kelp cultivation starts by collecting reproductive sporophytes from the wild and releasing zoospores that settle on seedspools (twine wrapped around PVC pipe) in the nursery (36). Zoospores then germinate into haploid gametophytes, which produce gametes after a few weeks that cross-fertilize and develop into diploid juvenile sporophytes. Once the sporophytes are 2-10 mm in length, they are outplanted to open ocean farm sites and grow to full-size (∼2-3 meters in length) sporophytes for harvest (36,37). In the nursery stage, growers have the most control over growing conditions and the microbial environment. Standard industry protocols employ strategies to reduce microbial growth and biofouling, such as autoclaving, pasteurization, and seawater filtration, along with physical removal of visible fouling and use of iodine solutions or weak sodium hypochlorite (36,38,39). However, these methods are costly, especially for small farmers (15)(15), and reducing the microbial load does not always effectively control pathogens and fouling organisms (40). Additionally, associations with bacteria are crucial for proper development and morphogenesis of macroalgae (41). For example, axenic cultures of the green alga *Ulva mutabilis* and brown alga *Ectocarpus sp.* show abnormal morphology (33,42), while co-cultures with microbial inoculants were phenotypically rescued (32). Similarly, the kelp *Laminaria* showed abnormal growth and died before reaching maturity in axenic culture (43). In the giant kelp *Macrocystis* the microbial community influences gametophyte development (44) and the presence of certain microbial taxa on gametophytes predicted higher sporophyte biomass (45). The removal of microbes in aquaculture is a standard practice to limit disease and biofouling pressure (36). However, losing beneficial microbes may have overlooked consequences. As such, microbial manipulation at the nursery stage of kelp cultivation may provide a unique opportunity to enhance productivity in kelp aquaculture by increasing the abundance of beneficial microbes while still limiting disease and fouling pressure (35).

The diversity of microbes associated with macroalgal hosts and the variability of these communities over space, time, and host condition make it challenging to determine which bacteria to target for microbial manipulations (46–48). Despite this variability, the microbiome of kelps appears to have stable taxa across space and time (49,50), making the core microbes promising targets for manipulation. There is a general assumption in the literature that bacterial taxa more consistently associated with host organisms (the core bacteria) are more likely to influence host function (51–53), making core microbes a good shortlist of candidate microbes to test for their ability to improve host health and performance. However, this prediction is largely untested and is based on often implicit assumptions. Here, we test the hypothesis that the core microbes are more likely to influence host function (herein termed the CIHF – Core Influence Host Function – hypothesis). Discussions of the core can be contentious as the thresholds used to define core taxa vary widely in the literature, from 100% to 50% frequency, sometimes alone and sometimes in combination with relative abundance or enrichment criteria (54–56). We argue that specific thresholds are less important than the concept and the ‘correct’ thresholds likely vary by study organism. We define core taxa as those that are consistently associated with a host and enriched on a host in comparison to the environment. Together, these criteria select for microbes that preferentially associate with a host by some repeatable mechanism (e.g. microbes attracted to host signals or food source, that resist or some other mechanism) and requiring enrichment excludes microbes that are common on a host because they are common in the environment (i.e. the common core of Risely, 2020). These criteria do not automatically exclude microbes with negative effects.

The CIHF hypothesis, though intuitive, remains largely untested. To address this gap, we assembled a broadly sampled 16S rRNA gene dataset surveying the microbiome *Saccharina latissima* in comparison to the surrounding environment over space and time from sites near Vancouver, Canada. We used this dataset to establish the distribution of bacterial taxa on *Saccharina* and define the core microbiome. We then assessed the distribution of these core taxa on cultivated kelp, both in nurseries and open ocean farms in British Columbia. In parallel, we cultured bacterial isolates from the surface of wild *Saccharina* and individually co-cultured 101 isolates representing 27 bacterial genera with *Saccharina* spores and assessed their influence on *Saccharina* development from spores to sporophytes in the lab. These genera vary in the strength of their association with *Saccharina* in the wild, enabling us to test the prediction of the CIHF hypothesis that more tightly associated isolates are more likely to increase host development rates. This work also serves as a proof of concept of the feasibility of using microbial manipulations as a tool in aquaculture.

## RESULTS

### Distribution of bacterial taxa on wild *Saccharina*

We compiled 16S amplicon data to establish the distribution of microbial communities associated with *Saccharina* across time and space in British Columbia, Canada and identify the core bacteria that are consistently present and enriched on *Saccharina* compared to the surrounding environment. We include a dataset originally assembled to investigate changes in the *Saccharina* microbiome during the spring freshet of the Fraser River that includes biweekly samples from April - July 2021 at four sites near Vancouver, Canada (48). We complemented this dataset with monthly samples over one year at one of these sites to capture microbiome variation through the winter. Overall, the microbial communities on *Saccharina* are distinct from those on rocks and in the water column (PERMANOVA pseudo-F_(2:329)_=40.072, R^2^=0.19588, p=0.0001; Fig. 2A). The microbiome on wild *Saccharina* varies seasonally and by site (two factor PERMANOVA site: pseudo-F_(3:181)_= 2.1350, R² = 0.02546, p = 0.0006; month: pseudo-F_(10:181)_= 5.8132, R² = 0.23104, p = 0.0001). Yet, a large portion of the microbiome is stable over time and space as visualized by plotting the relative abundance of the most common genera [Fig. 2C]. This broad dataset enables us to identify the core taxa that are stably and specifically associated with *Saccharina* across sites and seasons.

**Fig. 1.**
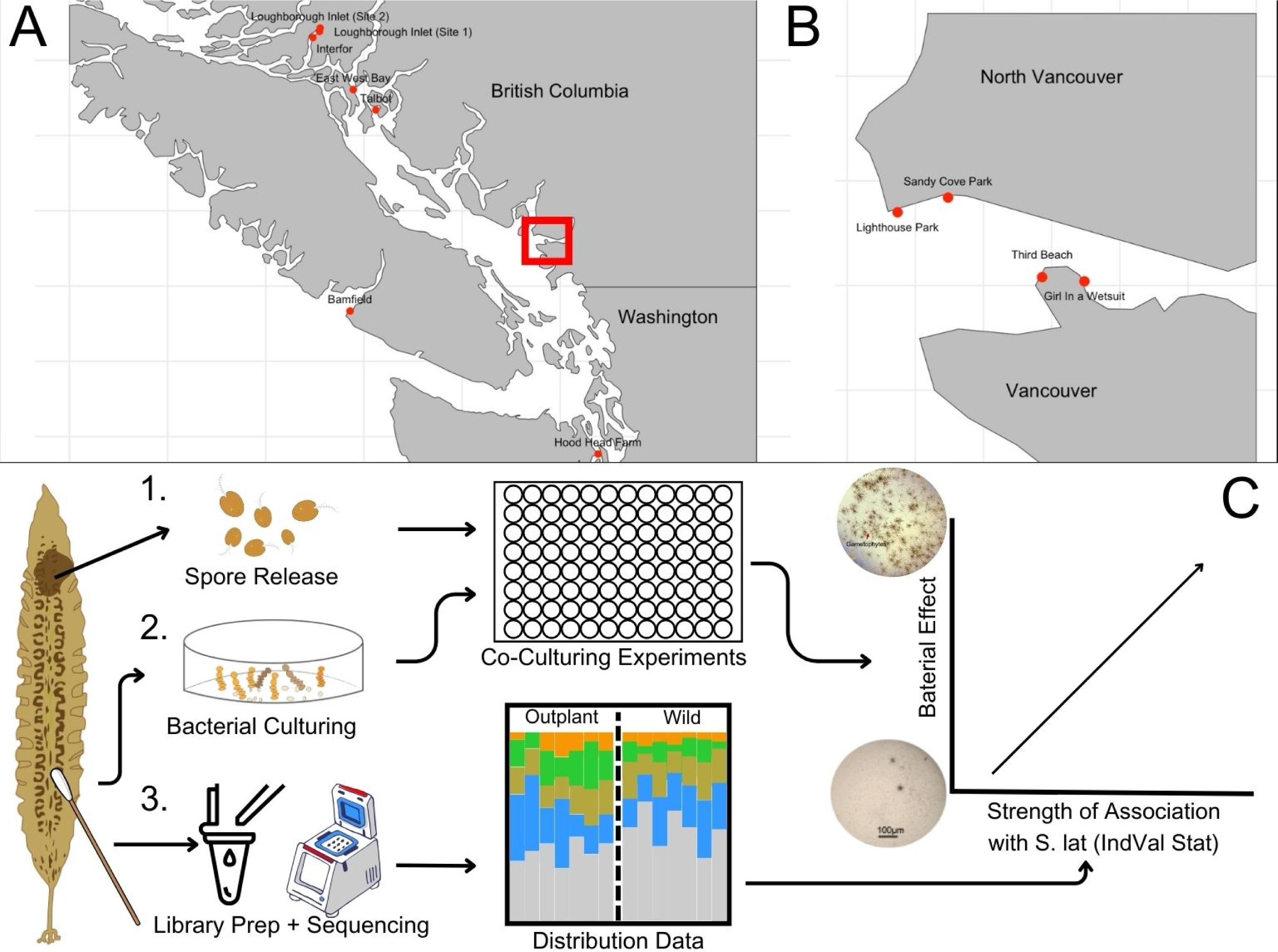
The experimental design for bacterial sampling and co-culturing *Saccharina* with bacterial isolates. **A.** Bacterial communities associated with *Saccharina* were sampled from various out-planted sites across British Columbia and northern Washington. Samples from Loughborough Inlet site 1 and 2 (LI), East West Bay (EWB), Talbot (TAL), Interfor (INT) and Hood Head Farm (HHF) were taken by Davis et al., 2023. Additional samples from Bamfield (BAM) were taken were taken for this study. **B**. Bacterial communities associated with wild *Saccharina* were sampled biweekly from April-July 2021 at various sites across Vancouver, BC. Additional samples were taken monthly from the Girl in a Wetsuit (GWS) site from July 2020-July 2021. Bacterial isolates used in part C were collected from GWS. **C**. 1. *Saccharina* zoospores were released and cultured with purified bacterial isolates (2) in the laboratory to assess the impact of different bacterial isolates on the early development of *Saccharina*. 3. Bacterial communities associated with *Saccharina* were analyzed using bioinformatics techniques to identify core bacterial taxa and quantify the strength of bacterial associations with *Saccharina* using IndVal analysis. Graphic made using Canva.

**Fig. 2.**
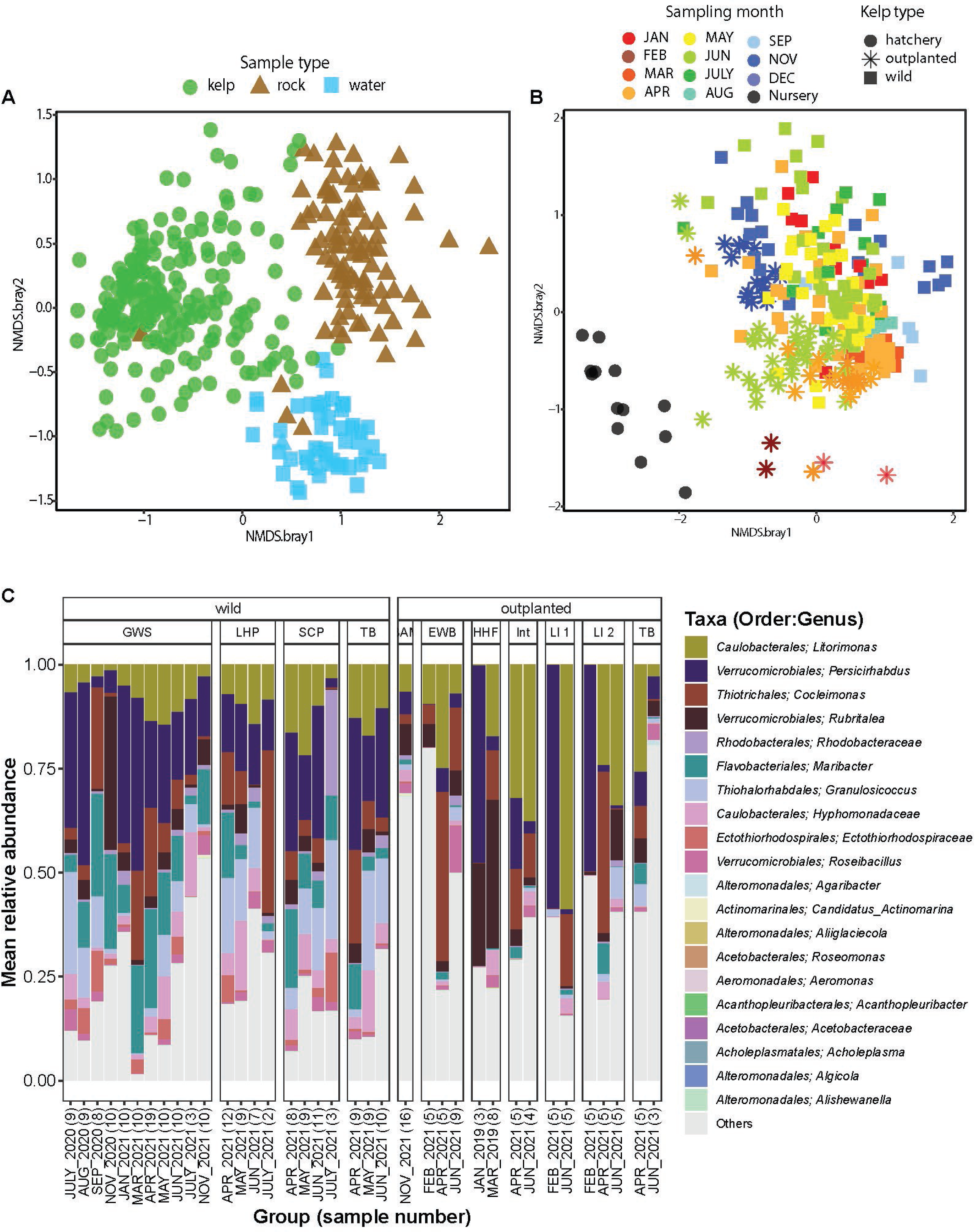
NMDS plots using Bray-Curtis dissimilarities and a taxa-plot were made to compare the microbiota found on different *Saccharina* populations and on surrounding abiotic substrates. **A.** Bacterial community composition on *Saccharina* compared to rocky substrate and seawater in the surrounding environment. **B.** Bacterial community composition on *Saccharina* samples collected from wild populations through the year in comparison to samples from cultivated *Saccharina* in the nursery and ocean farm sites. **C**. Most abundant taxa found on several outplanted and wild *Saccharina* populations. Abundances are summed across all samples collected at the same timepoint for a given site.

### Core bacteria of wild *Saccharina*

We used indicator species analysis (IndVal) to identify core taxa associated with wild *Saccharina* at the genus and ASV level [Table S2 and S3]. We set a threshold of >0.7 for the IndVal index because this selected only taxa that were present on the majority of samples and enriched on *Saccharina* compared to the environment, consistent with our definition of the core. This resulted in 8 core bacterial genera and 12 core bacterial ASVs. The same core taxa were identified when restricting to equal sample size per site per month [Table S2]. The core bacterial genera were *Persicirhabdus*, *Maribacter*, *Litorimonas*, *Granulosicoccus*, *Hellea*, *Cocleimonas*, *Ectothiorhodospiraceae* (genus unidentified) and *Marixanthomonas*. Additionally, there were three core ASVs belonging to *Rubritalea*, *Blastopirellula* and *Loktanella,* which were not core at the genus level [Table S2 and S3].

We plotted the frequency and mean relative abundance of the 12 core bacterial ASVs on wild *Saccharina* across the four sites and over time [Fig. 3]. We also plotted the sum of remaining ASVs within the 8 core bacterial genera to visualize any tradeoffs within genera across sites or over time. In most core genera, such as *Cocleimonas*, *Hellea*, and *Maribacter*, the core ASV is predominant. *Persicirhabdus* may tradeoff seasonally as a second ASV in winter samples [Fig. 3], though winter sampling is too limited to draw conclusions. The dynamics within *Granulosicoccus* differ, with multiple lower prevalence ASVs contributing to very high prevalence at the genus level (90%) [Fig. 3].

**Fig. 3.**
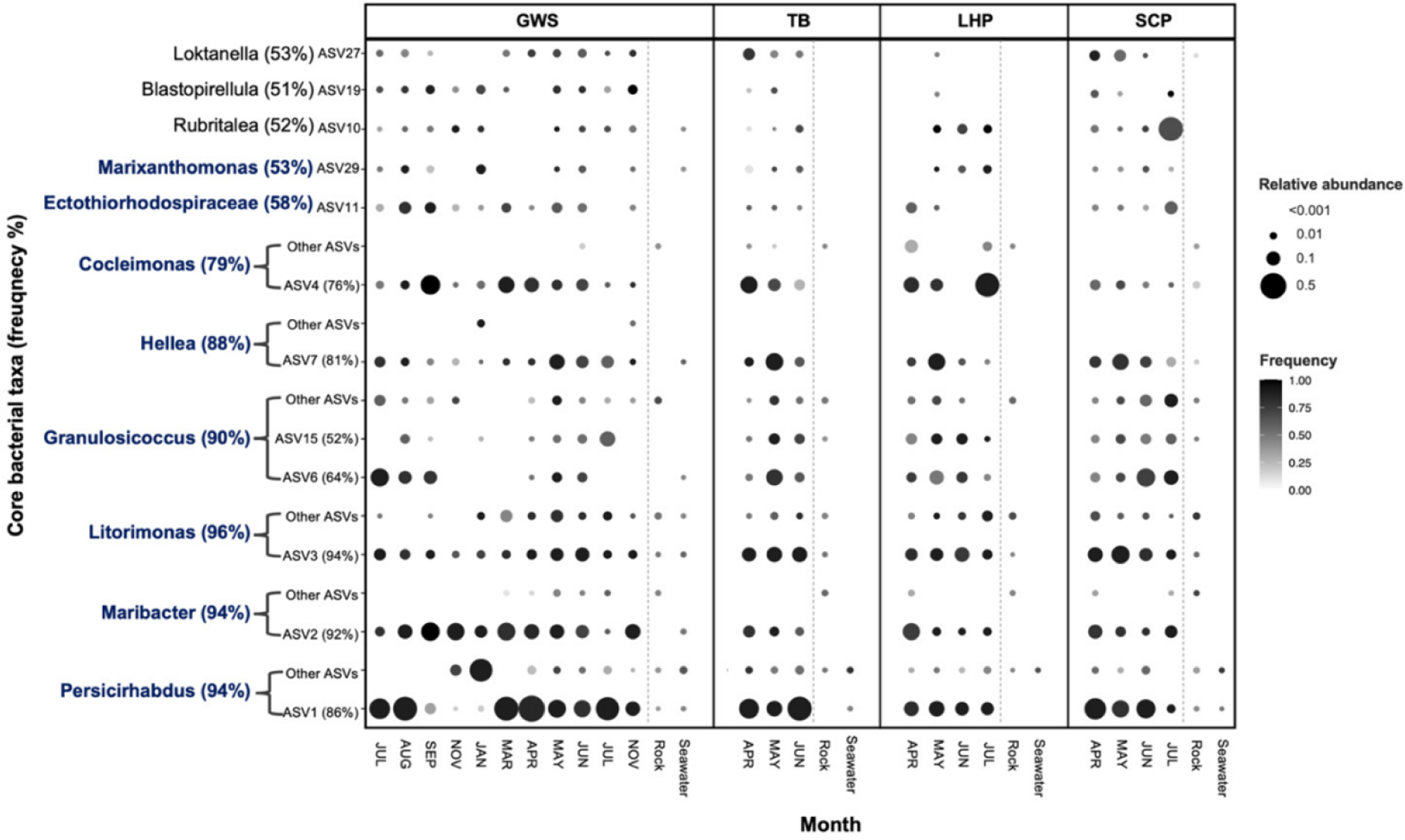
Distribution of core taxa S. latissima in four intertidal sites (GWS, TB, LHP and SCP). The dots represent the average relative abundance and frequency of core bacterial taxa across four locations over time. The y-axis displays the core bacterial taxa identified at the genus level in blue text and at the ASV level in black text, along with the overall frequency percentage of the core bacteria. The "other ASVs" refers to summed abundance of all non-core ASVs assigned to the core bacterial genus. The monthly seawater and rock samples data were aggregated across all dates.

### Microbiome on cultivated versus wild kelp

We compared microbiome composition and the distribution of core bacteria between wild and cultivated kelp by reanalyzing a previously published dataset of cultivated *Saccharina* sampled across a developmental time series (57). Juvenile sporophytes (∼1-2 cm in length) were sampled from cultivation seedspools in one nursery on Vancouver Island, BC and one nursery in Washington. Juvenile sporophytes on the seedspools were then outplanted to seven ocean farm sites (hereafter referred to as farms; Fig. 1) and sampled two, four, and six months after outplanting; at six months they were mature sporophytes (57). The microbiome composition is significantly different in the nursery, on outplanted cultivated kelp, and on wild kelp (Fig 2B; PERMANOVA: pseudo-F_(2:293)_= 16.866, R² = 0.10324, p = 0.0001; pairwise PERMANOVA p<0.001) for all comparisons. The comparison between wild and cultivated kelp is confounded by site differences and partially confounded with seasonality and sampling method. We ran two additional analyses to better account for the differences in seasonality and sampling method between wild kelp and some of the cultivated kelp samples. First, we restricted the comparison to BC kelp sampled at the same time of year (April, June, and November/December) and ran a nested PERMANOVA comparing cultivated and wild nested within month (pseudo-F_(5:163)_= 8.9969, R² = 0.21629, p = 0.0001). Second, we restricted the comparison to BC samples collected in June and compared wild kelp that were sampled by swabbing to cultivated outplants sampled by the same swabbing protocol and to cultivated kelp sampled from whole tissue (PERMANOVA: pseudo-F_(2:76)_= 5.977, R² = 0.13591, p = 0.0001). Follow up pairwise PERMANOVA from only June samples comparing wild, cultivated swab, and cultivated whole tissue show that samples taken by swab and from whole tissues differ (outplant swab versus outplant tissue: R² = 0.06, p = 0.003), but wild vs cultivated explains a larger portion of varation (swab wild versus swab outplant: R² = 0.11, p = 0.001); swab wild versus tissue outplant: R² = 0.12, p = 0.001). In all cases, wild and cultivated kelp microbiome communities differ, though we cannot rule out site as a cause of this difference.

We then assessed the distribution of *Saccharina* core taxa on cultivated *Saccharina* in nursery facilities and farms [Fig. 4]. These core taxa were absent on *Saccharina* seed spools at the nursery facilities, and all except *Persicirhabdus* were absent two months after outplanting to ocean farm sites [Fig. 4]. Interestingly, most of the core bacterial taxa colonized *Saccharina* gradually over time at the six open ocean farms originally studied by Davis et al. (2023) and sampled between 2 and 6 months after outplanting [Fig. 4]. This pattern is reinforced by the observation that all core bacteria from wild kelp were found on the cultivated kelp at the Bamfield site that had been outplanted for 12 months [Fig. 4]. This suggests that the core bacteria identified in wild kelp populations are recruited to cultivated kelp over time.

**Fig. 4.**
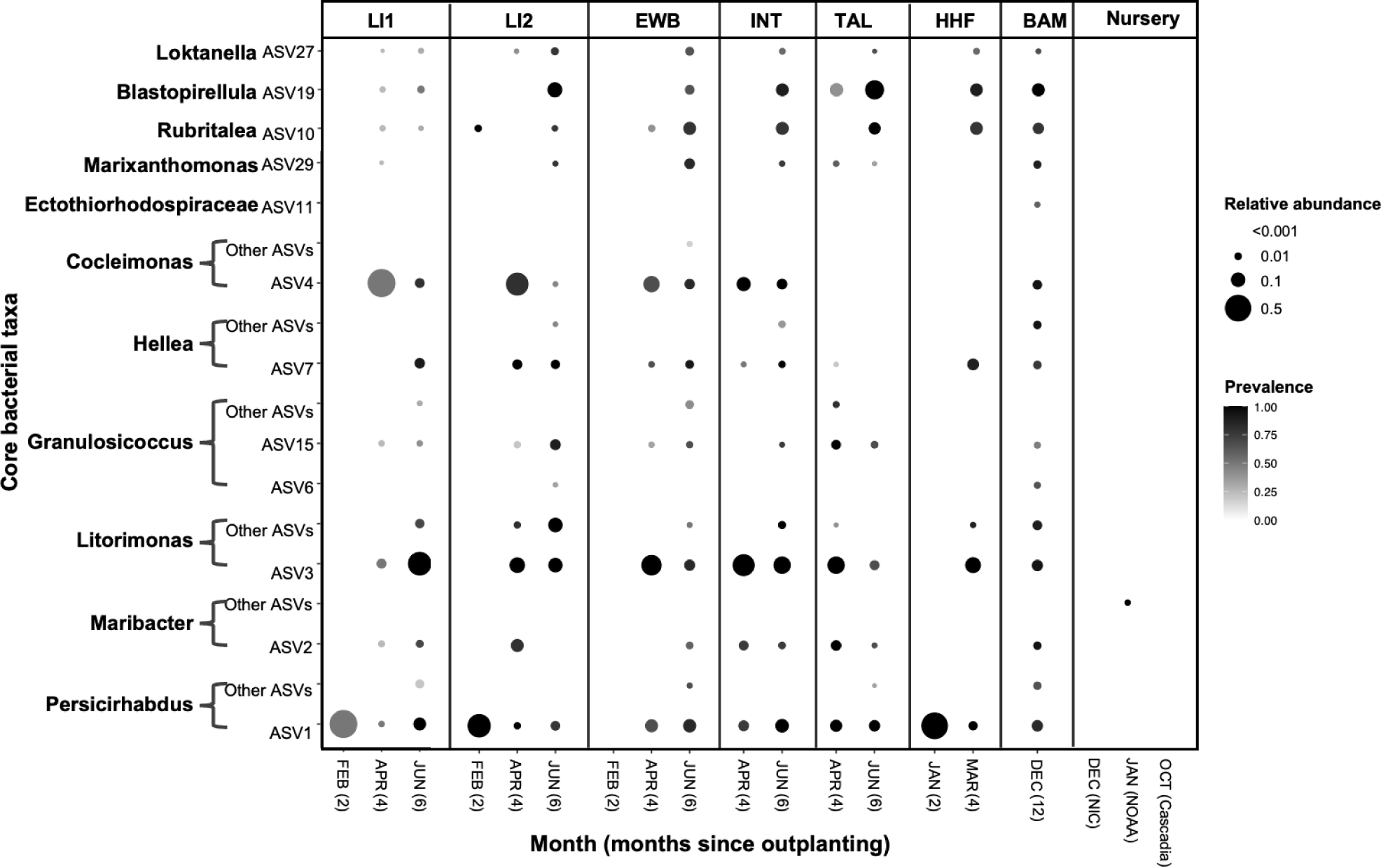
The average relative abundance and frequency of *Saccharina* core bacterial taxa identified from wild populations (shown in Fig. 3) on cultivated kelp. Cultivated kelp samples come from three nurseries and seven ocean farm sites. Ocean farm sites except BAM were sampled at two, four, and six months after outplanting, as described in Davis et al., 2023. Two month samples failed for all INT and TAL sites, and all but 1 sample for LI2 and EWB. See Fig. 3 for additional notes.

### Cultured bacterial isolates

We cultured bacteria from the surface of meristem tissue of wild *Saccharina* from the “Girl in a wetsuit” site. In an effort to culture as wide an array of bacteria as possible, we used a total of seven types of culture media (1. Full-strength Difco Marine, 2. Half-strength Difco Marine, 3. One-fourth strength Difco Marine, 4. One-tenth strength Difco Marine, 5. Full-strength Difco Marine with alginate acid, 6. Full-strength Difco Marine with antibiotics, and 7. Full-strength Difco Marine with ground kelp tissue), and isolated bacteria at eight timepoints from August 2020 to February 2022 [Table S1]. We obtained 127 bacterial isolates and identified them through Sanger sequencing of the 16S rRNA gene; 27 out of 127 failed to sequence [Table S1]. The isolates belong to 32 genera. Overall, we cultured 5.9% of the bacterial genera detected in the wild *Saccharina* microbiome by 16S rRNA Illumina sequencing (32 out of 535 genera detected). Three cultured genera represented bacterial genera identified as core: *Maribacter, Hellea* and *Litorimonas*. Based on phylogenetic analyses, the isolates representing the core genera were distinct strains compared to the core ASVs. *Maribacter* isolate M16 matched ASV14, not core ASV2 [Fig. S2]. *Litorimonas* isolate H37A matched ASV18 rather than core ASV3 [Fig. S1]. In addition, we found that core ASV7 within the *Hyphomonadaceae* belongs to the genus *Hellea* [Fig. S1]. We tested 101 of these bacterial isolates in co-culture experiments with *Saccharina* and assessed the ability of 99 isolates to produce auxin.

### The effects of bacterial inoculation on *Saccharina* cultures

We conducted four trials where we co-cultured *Saccharina* zoospores with bacterial isolates to observe their effect on gametophyte germination and the number of sporophytes produced. The number of isolates tested and number of replicates differed across trials (trial 1: 62 isolates, n=2 per isolate; trials 2 and 3: 25 isolates, n=4 per isolate; trial 4: 11 isolates, n=8 per isolate). Most isolates were tested only once (94 isolates), while seven were tested in multiple trials. The influence of bacterial isolates on gametophyte germination and number of sporophytes was analyzed separately to account for variation across trials, but are plotted together for simplicity [Fig. 5]. Within each trial, the effect of bacterial isolates on kelp was calculated with t-tests comparing the log_2_fold change of gametophyte coverage and number of sporophytes produced for each bacterial isolate to the control treatment (*Saccharina* spores without bacterial inoculants), followed by Benjamini-Hochberg correction for multiple comparisons [Table S4].

**Fig. 5.**
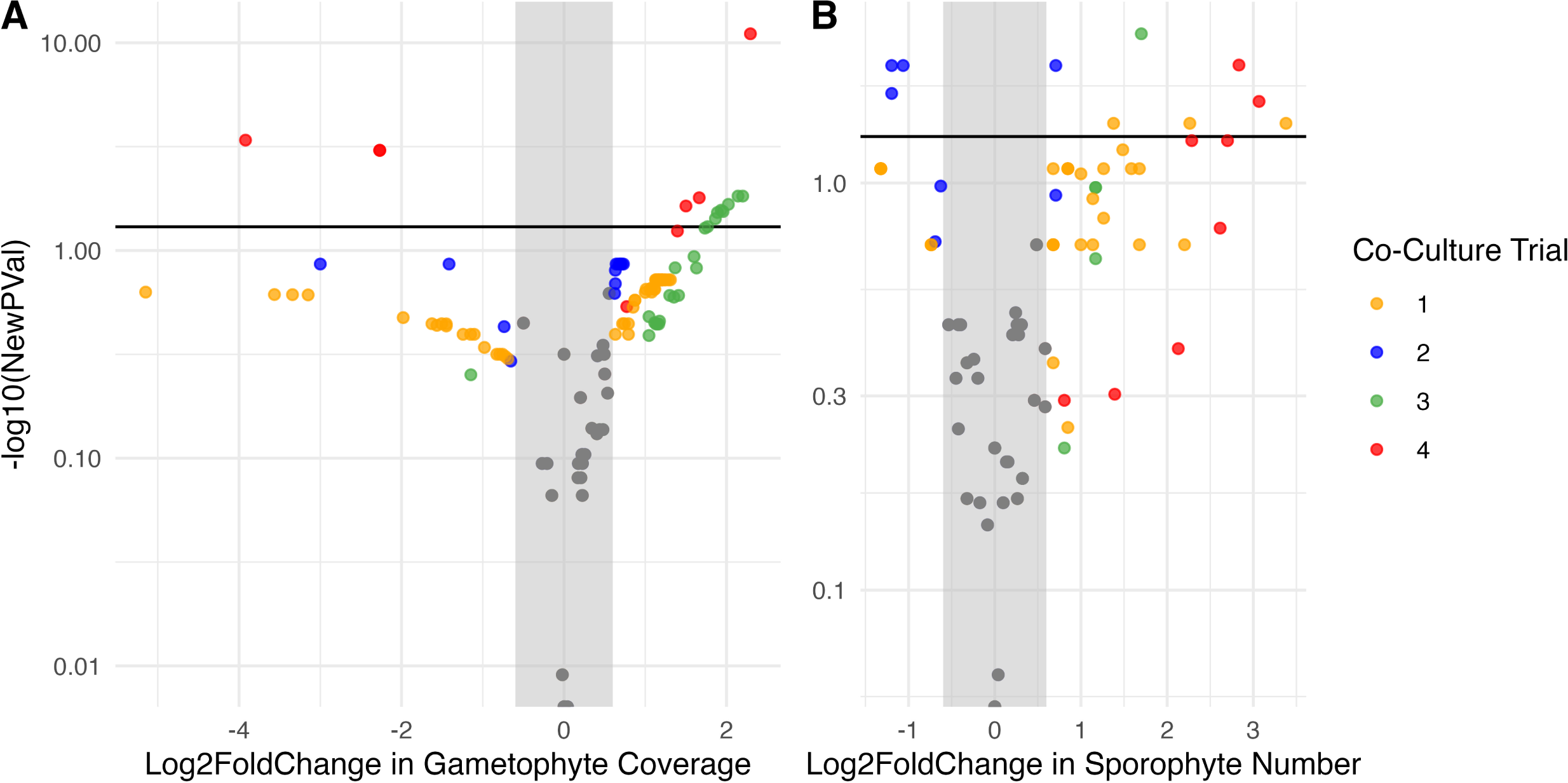
Volcano plot depicting the effect of bacterial inoculants on *Saccharina* across all four trials. The x-axis depicts the log2 fold change for each bacterial isolate compared to the control groups in each experimental trial on (A) gametophyte percent cover and (B) the number of sporophytes produced. The y-axis displays log10 Bonferroni-corrected p-value from two-tailed t-tests comparing the control group to co-culture treatments with each bacterial isolate. Isolates are colored by co-culture trial, and grey if their effect was not significant.

Across all trials, 33 isolates (28 of which were unique) had a significant positive impact on gametophyte coverage, while 5 isolates (4 of which were unique) had a significant negative effect [Fig. 5A]. 23 isolates (17 of which were unique) had a positive impact on sporophyte number and 4 unique isolates had a significant negative effect [Fig. 5B].

In trial 4 only, we assessed the persistence of the inoculated bacterium and the overall microbiome in culture wells using 16S rRNA sequencing. We confirmed that four of the seven inoculated bacterial isolates (ASV14.*Maribacter*, ASV12.*Sulfitobacter*, ASV18.*Litorimonas*, ASV5.*Pseudoalteromonas*) were indeed overrepresented over time in the kelp culture trial 4 [Fig. S3].

To assess the hypothesis that the core bacteria are more likely to influence the growth and development of kelp, we asked whether the ecological distribution of bacterial genera in the kelp microbiome predicted their effects on *Saccharina* development in co-culture using linear mixed effects models with trial as a random effect. There is a significant relationship between the strength of association between bacterial genera with *Saccharina* (measured by IndVal value at the genus level) and the effect of bacterial isolates on number of sporophytes produced (p = 0.00051, F = 12.957) [Fig. 6A] and gametophyte coverage (p = 0.0167, F = 5.937) [Fig 6B].

**Fig. 6.**
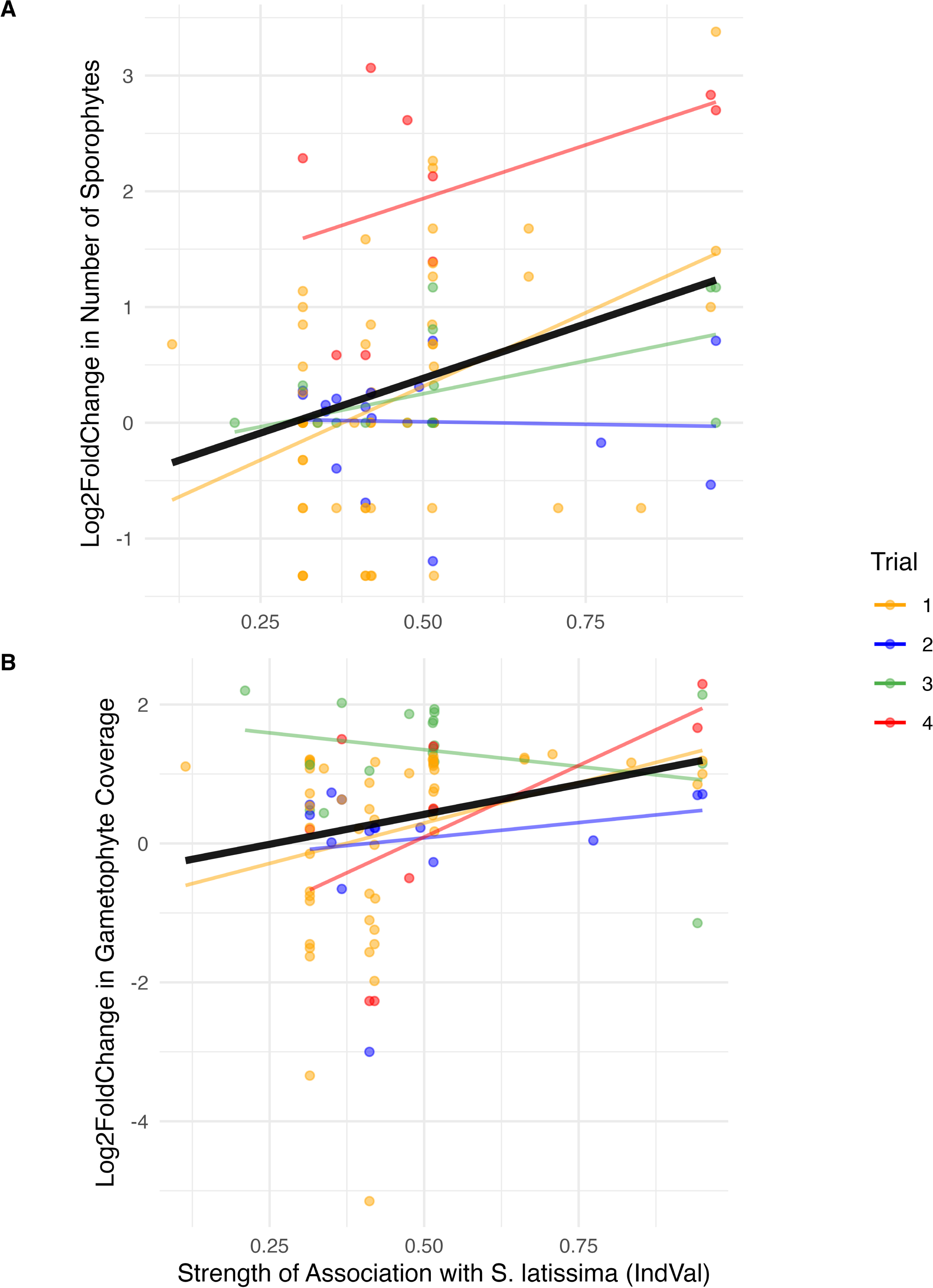
The relationship between strength of association with wild Saccharina (IndVal statitistic) and effects of bacterial inoculation on *Saccharina* development. The y-axis displays the log2 fold changes for each bacterial isolate compared to the control groups in each experimental trial, for (A) gametophyte coverage % and (B) the number of sporophytes. Points and trend lines are colored co-culture trial, while black trend-lines represent the relationship across all trials.

Overall, these findings suggest that bacterial genera more strongly associated with *Saccharina*are more likely to increase gametophyte grown and induce the production of more sporophytes.

We then asked if the variation in this relationship was explained in part by the ability of bacterial isolates to produce the growth hormone IAA (auxin) as measured by the Sokolov colorimetric protocol using the Salkowski Reagent (58). Of the 129 isolates plated from glycerol stocks, 99 grew and were tested for IAA production; 59 of the 99 produced IAA in measurable quantities (Table S1). We treated IAA production as a yes/no value, as we did not standardize bacterial culture density at the time of the IAA assay. We found that bacteria that produced IAA were more strongly associated with *Saccharina* than those that did not produce IAA using a t-test comparing the IndVal stat of IAA producers and non-producers (t-test: p = 0.0002). We then asked whether IAA producers were more likely to influence kelp growth and development in co-culture. We restricted this analysis to co-culture trial 1 where the most isolates were tested to maximize our power to detect a result and used t-tests to compare the log_2_fold change of number of sporophytes and gametophyte coverage between IAA producers and non-IAA producers. There was no difference (Sporophyte number: p = 0.5825; Gametophyte Coverage: p = 0.3052).

We asked whether the bacterial isolates tested repeatedly, and different isolates from the same genus, have a consistent influence on *Saccharina* germination and development. Results for the seven bacterial isolates tested three or more times varied across trials but the trends were generally consistent; no isolates switched from significant positive to significant negative effects or vice versa [isolates tested repeatedly are bold in Table S4]. Only one isolate, *Maribacter* M16, always significantly increased the number of *Saccharina* sporophytes produced, while inoculating the remaining isolates yielded significant changes in gametophyte coverage or sporophyte numbers in some but not all trials [Table S4]. We observed more variation across unique isolates from the same genus. The highly culturable genera *Pseudoalteromonas*, *Shewanella*, *Sulfitobacter*, and *Vibrio* each had isolates that induced significant increases and others that induced significant decreases in the number of sporophytes [Table S4]. We note that culturing biases strongly influence the pool of cultured isolates such that some genera are represented by many isolates.

## DISCUSSION

We investigated the distribution of bacteria on wild and cultivated *Saccharina,* and the effect of bacterial isolates on *Saccharina* growth and development in co-culture experiments. Together, these data enabled us to investigate the relationship between microbial distribution and functional impact, one of the first studies to empirically test this hypothesized relationship (52,53,55). We found distinct bacterial communities associated with *Saccharina* compared to the surrounding environments and identified a strong and consistent group of core taxa across four sample sites near Vancouver, Canada [Fig. 3]. Additionally, our co-culture experiments showed that bacteria can significantly influence kelp sporophyte development [Fig. 5], and that there is a significant, positive relationship between the strength of association of microbes on *Saccharina* and the functional effect of these microbes on host development [Fig. 6]. These findings support the CIHF hypothesis, which predicts that the most prevalent and enriched microbes are more likely to influence host function.

The rationale of the CIHF hypothesis is rooted in the assumption that 1) consistent ecological associations between a host and microbe are observed when there are repeatable mechanisms (e.g. microbes attracted to food source, host attracts particular microbes, vertical transmission) that lead to the association re-forming, and 2) that the existence of consistent associations increases the likelihood of hosts and/or microbes evolving adaptations that promote the association and/or increase the reliance on [by]products of the partner. Importantly, consistent associations can lead to evolution of dependence even in the absence of strict or obligate relationships, giving rise to the functional redundancy often observed in symbiotic interactions. For instance, the bacterium *Roseobacter* sp. is attracted to *Ulva* sp. by DMSP, and produces morphogens crucial for *Ulva*’s development, but many bacteria can substitute (59,60). The CIFH hypothesis applies to all associations that reform repeatedly, not only those that benefit the host. Although core taxa were absent from cultivated *Saccharina* in nurseries, they colonized outplanted kelp after a few months [Fig. 4], suggesting these repeatable associations are the result of environmental acquisition and likely involve a combination of microbial attraction and host filtering mechanisms. Their repeated presence suggests that core taxa have co-evolved to some extent to form consistent associations and further studies needed to understand the underlying mechanisms.

The relationship between microbial taxonomy and function is complex due to functional redundancy (61) and variation in function observed even between different strains of the same species (62,63). As a result, the functional influence of core microbes on a host cannot be determined from taxonomy alone. In our co-culture trials, we observed several instances of contrasting effects for isolates from the same genus, including *Sulfitobacter* and *Pseudoalteromonas* [Fig. 5], and even contrasting influence of strains within the core genus *Maribacter* [Table S4]. Yet, taxonomic information is still a predictor of function though the level of taxonomic conservation of functional traits varies widely across functions (64) and some taxa, such as *Pseudoalteromonas* and *Vibrio,* are more functionally variable than others (63,65). Here, we found a significant relationship between the distribution of bacterial genera on *Saccharina* and the influence of isolates within the genus on *Saccharina* development.

Importantly, this relationship was observed at the genus level as we were unable to culture the core ASVs to test their influence in the lab. This suggests that knowing which taxa are present in this system, in addition to the functions they may perform, is valuable and motivates future investigations of function and taxonomy in tandem.

We explicitly tested the relationship between distribution and one function thought to influence algal growth: production of the phytohormone auxin (IAA) by bacterial isolates. Auxins mediate plant-microbe interactions, both in mutualisms and parasitism, and are an interdomain signaling molecule (66,67). Auxin production and signaling is not limited to land plants; there is evidence of auxin influencing growth of green (68), red (69), and brown (70) algae, and auxin is produced by a wide range of algae and bacteria, making it a plausible candidate for mediating kelp-microbe interactions. We found a high degree of functional redundancy in that more than half of the isolates tested produced auxin in the form of IAA, though there was no relationship between an isolate’s ability to produce auxin and its effect on *Saccharina* in co-culture [Fig. 7]. However, isolates that produce auxin belong to genera that are more strongly associated with *Saccharina* [Fig. 7], suggesting that bacteria with the ability to manipulate host growth may preferentially associate with *Saccharina*. It should be noted that auxins and indole compounds produced by marine bacteria are also known to mediate both intra and inter-species interactions such as biofilm formation and quorum sensing (71,72), so further experiments are needed to determine the mechanisms that underlie the observed relationship.

**Fig. 7.**
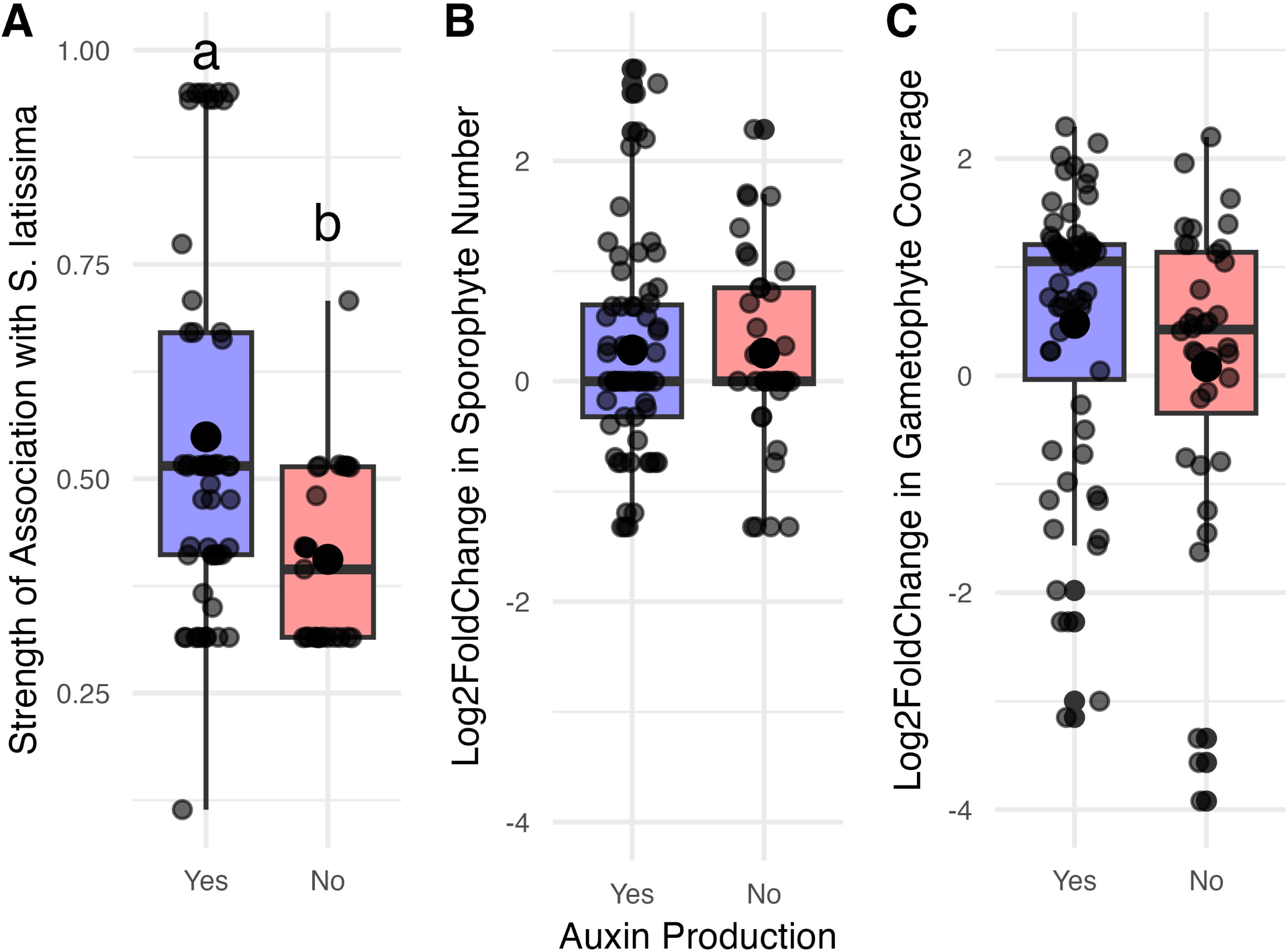
The relationship between isolate auxin production and distriubtion and effect in co-culture. The x-axis shows whether auxin was produced in a detectable amount in IAA-screening experiments. The y-axis shows (A) the strength of an isolate’s association with Saccharina (IndVal stat), and the isolates effect in co-culture on number of sporophytes produce (B) or change in gametophyte coverage (C). Significant difference between group averages as determined by a t-test are noted by letter.

Our results suggest that bacterial manipulation may be useful to increase productivity in an aquaculture setting by facilitating gametophyte settlement and development of sporophytes in commercial kelp at nursery stages. We found that several bacterial isolates significantly influenced both spore settlement/gametophyte germination and sporophyte development of *Saccharina* in co-culture [Fig. 5]. While there is variation across trials, we also observe broad consistency in the effect of individual isolates that were tested in multiple trials despite variation across trials and no instances of flips between significant positive and significant negative effects [Table S3]. This study inoculated one bacterial strain into an existing consortium because the kelp could not be cultured axenically; this is a limitation of the study that means we cannot firmly establish causality. This is also a strength of the study because it shows that inoculation of isolates repeatedly influences host function in a conditions that mimic the non-axenic conditions of kelp nursery facilities. There are abundant avenues to test strategies that might result in increased host function, such as inoculating multiple bacterial strains together and manipulating the background consortia through antibiotics. We focused our manipulation trials on the gametophyte and juvenile sporophyte stages of development because this stage, grown in aquaria in kelp nurseries, is the easiest for growers to manipulate. Altering growth and development in the nursery could potentially benefit kelp aquaculture productivity directly by decreasing the time spent in the nursery stage, or by increasing sporophyte yield at harvest. Increasing growth and development rates in the nursery was identified as the stage with the greatest potential to reduce cultivation costs and enhance productivity, thereby increasing the economic viability of kelp aquaculture in North America (10). Furthermore, the presence of some microbial taxa on Macrocystis gametophytes was correlated with sporophyte yield at harvest (45) Overall, manipulating the microbiome of early kelp development stages has great potential to influence harvest yields in kelp aquaculture. There are abundant avenues for further experimental testing, including testing consortia of microbes rather than single strains.

One of the challenges to successful microbial manipulations is the ability of the inoculant to colonize within a complex community (73,74). While we followed protocols from industry for spore release and cultivation designed to minimize the presence of microbes (37,39), these protocols do not completely remove microbes. Consequently, inoculated bacteria were added to a complex community, yet most were detected in culture wells 28 days after inoculation, suggesting that they can persist early in the development process despite not being detected in nursery samples [Fig. S3]. The ability of inoculants to persist in the existing microbial community is crucial given that axenic kelp cultures often fail (43). Many of the successful inoculants also had significant effects on both gametophyte coverage and sporophyte development in co-culture [Fig. 5]. Further investigations are needed to understand the competition and colonization dynamics of inoculated bacteria.

## CONCLUSION

This study tests the CIFH hypothesis, and whether this framework can be useful to guide microbial manipulation efforts for kelp aquaculture. We found a significant positive correlation between a microbe’s frequency and enrichment on wild *Saccharina* and its effect on host development in co-culture trials, indicating that core microbes are more likely to influence host function. Furthermore, core microbes colonized cultivated kelp over time after outplanting to ocean farm sites, suggesting the presence of horizontal transmission mechanisms that sustain specific microbe-kelp associations. Our co-culture experiments demonstrate the potential for microbial inoculations to significantly influence host development in a consistent manner.

Together, these results suggest that a taxonomic core framework can be used to identify microbial taxa more likely to influence host development, and that these microbes can be successfully inoculated and affect the early development of *Saccharina*, laying the foundation for developing microbial tools to enhance kelp aquaculture.

## MATERIALS AND METHODS

### Sample collection

*Wild Saccharina spatial survey*: Microbial DNA samples were obtained from the surface of meristem tissue of *Saccharina* bi-weekly at four intertidal sites from April to July 2021: Girl in a wetsuit, GWS (49.302718, -123.126230), Lighthouse Park, LHP (49.329846, -123.264521), Sandy Cove Park, SCH (49.340294, -123.226243), and Third Beach, TB (49.302322, - 123.157299), as described in Schenk et al., in revision. Individual kelp were rinsed with 0.22 μm filtered sterile seawater for 10 seconds to remove loosely associated microbes and then swabbed with a Puritan® sterile swab for 15 seconds, covering an approximate area of ∼3x3 cm^2^. Swabs were immediately stored in 2 mL cryovials (VWR/Avantor company). Bare rock substrates were swabbed as a comparison to non-host associated microbial communities (three sample per site per timepoint). Three water column samples were also collected from adjacent seawater at each sampling site per visit to characterize microbial source pool communities by filtering seawater onto a 0.22um Millipore Sterivex™ unit. Samples were transported back to the lab and stored at -70 C within 4 hours of collection until DNA extraction.

*Wild Saccharina temporal survey*: Additional samples were taken monthly at one site (GWS) from July 2020 to July 2021 and in November 2021 for this study to survey the microbial community throughout the year. These samples were taken as above, except that live *Saccharina* specimens were placed in an icebox with local seawater and transported to the laboratory and swabbed in the laboratory within two hours of collection then placed in cryovials and stored at - 70°C until DNA extraction. Rock swabbing and water filtering took place in the field as described above.

*Cultivated Saccharina:* Microbiome samples of cultivated kelp were originally collected by Davis et al. (2023) from two nurseries (North Island College in BC, Canada, and NOAA in WA, USA) and six ocean farm sites [Fig. 1] and were used to assess microbial changes through the cultivation cycle. Here, we reanalyzed these data to assess the presence of a core microbiome over time at open farm sites. Samples from a third nursery (Cascadia Seaweed in BC) and one additional ocean farm site (Bamfield) were taken for this study. Samples in the WA and Cascadia nurseries were collected by swabbing a 3 cm^2^ area of seedspool when sporophytes were 1-4 cm in length. Additional samples were taken of aquarium tank walls by swab and aquarium water by filtering, as above. Nursery samples from NIC were collected by cutting 5 cm lengths of seedspool twine sporophytes that were 1-4 cm in length and extracting from whole tissue, as described in Davis et al. (2023).

Samples of outplanted kelp were collected roughly every 2 months after outplanting to ocean farm sites for all farm sites except Bamfield, which was sampled 12 months after outplanting only. Outplanted kelp were sampled by the swab protocol described above or from whole tissue as described by Davis et al., 2023. Samples from Bamfield, Washington, and for roughly half of the 6 month time point samples to other BC farm sites were swabbed. At the two, four and the other half of 6 month time point, outplanted cultivated kelp from BC farm sites were sampled by extracting microbial DNA from whole tissue due to pandemic travel restrictions.

Significant but small differences between the swab and whole tissue DNA extraction methods were previously detected (57). At the six and twelve month post-outplant time points, the *Saccharina* were mature sporophytes similar in size to the wild-collected sporophytes.

Additionally, we collected samples from nearby non-host substrates (unseeded cultivation ropes) except for the Bamfield site. All samples were stored at -70°C until DNA extraction.

### Library preparation for 16S rRNA amplicon sequencing

For the samples characterized in this study, DNA was extracted from swabs and Sterivex filters for 16S rRNA gene amplicon sequencing using DNeasy 96 PowerSoil Pro Kit (384) (QIAGEN), following the manufacturer’s recommended protocols. PCR amplification using 30 cycles for bacterial DNA targeted the V4 region of the 16S rRNA gene using primers, 515f: 5’– GTGYCAGCMGCCGCGGTAA–3’ and 806r: 5’ – GGACTACHVGGGTWTCTAAT –3’ (75) using dual-index barcoded primers (76). Then, PCR products were visualized by gel then quantified using Quant-IT Pico Green® dsDNA Assay Kit (Invitrogen), and pooled at equal concentration (10 ng) of each sample. The pooled library was purified using MoBio UltraClean PCR clean-up kit. Bioanalyzer trace was conducted at the BRC Sequencing Core, University of British Columbia to assess the quality and target size of the pooled libraries before sending the final samples to the Hakai Institute Marna Genome Lab, Quadra Island, BC for sequencing via the Illumina MiSeq Plantform (MiSeq Reagent Kit v3 (600cycle); Illumina Cat # MS-1023003). Library preparation was carried out using using the same steps for Schenk et al., in revision and Davis et al, 2023.

### Bioinformatics and statistical analysis of 16S libraries

Demultiplexed FASTQ files for new data as well as data from Schenk et al and Davis et al. (2023) were processed together. Merging sequences from multiple datasets, quality filtering, trimming, dereplication, chimera removal, inference of ASVs and taxonomic assignment against the SILVA 138 (77) database were processed with The Divisive Amplicon Denoising Algorithm2 (DADA2) pipeline (78) in R environment (version 3.6.1). DADA2 employs a model-based approach for correcting amplicon errors without constructing Operational Taxonomic Units (OTUs) (79). We adhered to the standard parameters outlined in the DADA2 pipeline tutorial. Specifically, we applied filtering with MaxEE=2, set trimming lengths to 240 and 160, performed chimera removal using consensus method, and assigned taxonomy using the naïve Bayesian classifier method. This taxonomic assignment was based on exact matching between ASVs and sequenced reference strains from the SILVA 138 database (77). In the process of filtering, we discarded ASVs if they accounted for less than 0.1% of total number of reads or were found in fewer than 5 samples. Additionally, chloroplast and mitochondrial DNA were removed for downstream analysis. Overall, we obtained 4,897 ASVs after filtering. The final products were then converted into phyloseq (version 1.28) format (80) in R for all the downstream analyses. We rarefied to 1500 reads per sample prior to beta-diversity analysis.

Non-rarefied data were used for taxa plots and IndVal analysis. We used nonmetric multidimensional scaling (NMDS) to visualize all samples included in this study based on Bray-Curtis dissimilarity (81). We conducted permutational analysis of variance (PERMANOVA) using adonis2 by margin and 9,999 permutations in the vegan package (version 2.5.7) (82) to test for differences among bacterial communities.

### Core bacteria of *Saccharina* at the genus and ASV level

To identify a suite of core bacterial taxa, we used indicator species (IndVal) (version 1.7.12) analyses using multipatt function within ‘indicspecies’ package (83) on all of the data from wild *Saccharina*. Analyses were run separately at the ASV and genus level. The IndVal metric is based on both specificity (a measure of the relative abundance of a bacterial taxon on *Saccharina* compared to environmental rock substrate and seawater samples) and frequency (a measure of its presence across *Saccharina* samples) of bacterial taxa (84,85). We defined the *Saccharina*-core bacterial taxa based on an IndVal value of a 0.7 or above, with a frequency of greater than 50%, meaning the taxon was present in more than half of the wild kelp samples. Our previous study comparing methods for identifying the core (86) led us to adopt this approach because we observed that the most consistent taxa were captured by this approach. The number of samples per site varies due to higher sampling intensity at the Girl in Wetsuit site, and because not all samples successfully amplified. We repeated the IndVal analysis after restricting to 10 samples per site per month to ensure that uneven sample size did not have a large influence on the composition of the core; the results were very similar. We visualized the distribution of the core bacteria by plotting the relative abundance and frequency across sampling sites over time in both wild and cultivated *Saccharina* using the ggplot2 package (version 3.3.6) (87) in R. Additionally, we used IndVal values of genera a measure of their ecological distribution and the strength of association with wild *Saccharina* and compared them to functional data for bacterial isolates in co-culture and their ability to produce IAA.

### Bacterial isolation

*Saccharina* tissues were collected from the “Girl in a Wetsuit” site, Stanley Park (49.302731694102754, -123.12599476839195) at 11 monthly low tide events in 2020 and 2021 and brought back to the lab for bacterial isolation within 2 hours. In the lab, meristems were rinsed with autoclaved natural seawater before swabbing with a Puritan® sterile swab for 15 seconds in the same manner of microbial DNA samples. Seawater was obtained from the Vancouver Aquarium, which has an intake pipe at 30 m depth in the Burrard inlet and is subsequently transported to the University of British Columbia by truck. Swabs were then streaked on different types of customized agar plates [Table S1] and incubated at 10°C for 4-10 days until bacterial colonies were visible. Aiming to culture a broad array of bacteria, we used seven types of culture media: 1. Full-strength Difco Marine agar, 2. Half-strength Difco Marine agar, 3. One-fourth strength Difco Marine agar, 4. One-tenth strength Difco Marine agar, 5. Full-strength Difco Marine with 1% of alginic acid (176.1g/mol)., 6. Full-strength Difco Marine with 1% of penicillin (100 mg/L) and 1% of streptomycin (25 mg/L), and 7. Full-strength Difco Marine with approximately 5 g (wet weight) of ground *Saccharina* tissue per liter. Plates were incubated at 10°C for 4-10 days until colonies were visible. Phenotypically distinct colonies were subcultured multiple times onto fresh standard (55.1g/L) Marine Agar Difco plates until pure cultures were obtained. All bacterial isolates obtained in this study were stored in 25% glycerol stock at -70°C and revived for cultivation for subsequent experiments.

### Identification of bacterial isolates

We identified bacterial isolates using Sanger sequencing of the 16S rRNA gene. DNA was extracted from bacterial colonies on Marine Agar Difco plates using the PrepMan Ultra Sample Preparation Reagent following the manufacturer’s standard protocols. The PCR amplification was performed with the thermocycler program: an initial DNA denaturation step at 94°C for 3 minutes, 30 cycles of DNA denaturation at 94°C for 1 minute, an annealing step at 55°C for 1 minute, an extension step at 72°C for 90 seconds, and a final extension step at 72°C for 7 minutes, using the bacterial universal primers, 515f: 5’– GTGYCAGCMGCCGCGGTAA– 3’ and 1492r: 5’ – TACGGYTACCTTGTTACGACTT –3’ (88). Final PCR products were sent for Sanger Sequencing at the Genome Quebec Innovation Centre, QC, Canada. Products were generally sequenced with both forward and reverse primers, but in some cases only forward (515F) primers yielded high quality sequence, reported in Table S1. All sequences were trimmed and quality filtered using Geneious Prime (version 2021.1) with a quality score cut-off of 20.

Sequences shorter than 200 base pairs were excluded. Taxonomic assignment at the genus level was carried out with the SINA aligner following standard settings (Pruesse, 2012, Bioinformatics, 28) against all databases within SINA: RDP (Ribosomal Database Project), LTP (The All-Species Living Tree Project), and SILVA (SILVA rRNA Database). Taxonomic assignment was generally The SILVA least common ancestor, except in cases where the LPT least common was more specific and LPT was used [Table S1].

### Phylogenetic analysis of isolates from core genera

We constructed phylogenetic trees to determine whether the bacterial isolates of *Hellea, Litorimonas* and *Maribacter* belong to the same clades as the *Saccharina* core ASVs. Sanger sequences for isolates, ASV sequences, and reference sequences of these bacterial genera from the SILVA 138 SSU database (77) and best BLAST (89) hits from NCBI were combined into FASTA files for the QIIME2 environment using the ShortRead and seqinr package in R. Sequences were aligned and the alignment masked to contain only alignment columns that are phylogenetically informative in q2-phylogeny pipeline in QIIME2 (90). Phylogenetic trees were then constructed by using RAxML rapid bootstrap method (replicates=100) with GTRCAT model. The resulting trees were annotated with information on isolation source from GenBank records when available. Phylogenetic tree visualization and annotations were performed in Interactive tree of life (iTOL) v4 (91).

### Co-cultivation of *Saccharina latissima* and isolated bacteria

*Saccharina latissima* spores were co-cultured with bacterial isolates to identify isolates that influence the biology of kelp and test the CIHF hypothesis. A total of 127 bacterial isolates were obtained [Table S1]. Out of these, 101 unique bacterial isolates that grew in liquid culture were tested in co-cultivation trials with kelp [Table S4]. Seven bacterial isolates were repeatedly tested to evaluate the reproducibility of their effects on gametophyte coverage and sporophyte production based on large positive *Maribacter* (M16) or negative *Vibrio* (2Man3) effects in trial 1 [Table S4] or because they are common associates of seaweeds (*Sulfitobacter* (Mal1 and MH2) *Litorimonas* (37A), *Rhodobacteraceae* (KM3), and *Pseudoalteromonas* (Man4).

*Cultivation of Saccharina:* We conducted four trials co-cultivating *Saccharina* with single bacterial isolates in Nov 2020, Jan 2021, April 2021, and Oct 2021. Each well of the 96 well plates were filled with kelp zoospores (preparation described below), a single bacterial isolate (preparation described below), and F/2 media to a final volume of 200 μL. Plates were incubated at 10°C under cool-white LED lamps (75 μmol photons m^-2^ s^-1^) with a 16h:8h light:dark cycle for 28 days. On day 14 of cultivation, water changes were implemented by carefully inverting the 96-well plate and refilling each well with fresh F/2 media. This process was designed to replenish the rich media, minimizing the potential effects of nutrient limitation. The F/2 media used throughout was autoclaved for sterilization.

For each of the four trials, we collected *Saccharina* blades with mature sorus patches from intertidal populations at the “Girl in a Wetsuit” site at low tide and transported them back to the lab for spore release within 2 hours in a cooler filled with unfiltered seawater collected at the site. We released spores according to published methods (36). Briefly, we excised mature sorus patches from the kelp blade with a sterile razor blade and rinsed the sorus tissue with autoclaved seawater. We then disinfected the sorus tissue with a 2% betadine solution for one minute, rinsed it with autoclaved seawater, and wrapped it in paper towels to dry overnight in the dark at 10°C. The next day, we submerged the dehydrated sorus tissue in autoclaved seawater for approximately 15 minutes or until the seawater turned brown and cloudy to induce spore release by osmotic shock. We counted the motile zoospores over a hemocytometer to calculate their concentration, and then plated approximately equal densities (∼5 motile zoospores/10 μL final volume) into each well of a 96-well culture plate; this was typically less than 5 μL zoospore solution and 195 μL F/2 media. We note that non-motile zoospores were also transferred to the 96-well plate and may have increased the viable cell counts.

To prepare the bacterial inoculum of each isolate we transferred a fresh colony from agar plates to liquid marine broth media (Difco) and incubated it without shaking in 13 mL screw cap tubes at room temperature for 72 hours. We then harvested the bacterial cells by centrifugation (3700 × g at 4°C for 10 min) and washed them three times with autoclaved F/2, then added 3mL of sterile F/2 medium to the pellet. We estimated the concentration of bacterial cells by measuring their optical density at 600 nm (92), then added bacterial inoculum from a single isolate to each of the culture wells, resulting in a final bacterial density of ∼1 × 10^7^ cells mL^-1^. The control group for each trial consisted of zoospores and F/2 medium without any added bacterial cells. The number of isolates and replicates vary across trials as follows: 62 isolates (n= 2) in trial 1, 25 isolates (n = 4) in trial 2, 25 isolates (n = 4) in trial 3, and 11 isolates (n = 8) in trial 4.

### Quantifying the impact of bacteria on *Saccharina*

To investigate the effect of bacterial isolates on the early life stages of *Saccharina*, we estimated the percent coverage of gametophytes and counted the number of sporophytes that developed. After 28 days of cultivation, we examined the entire 96-well plate under the inverted microscope. Gametophytes, characterized by their singular, elongated form with branching structures, can be easily distinguished from the developing multicellular blade structure of sporophytes by microscopy [Fig. S4]. We did not systematically record the timing of gametophyte germination or sporophyte appearance. However, we observed that germination to gametophytes typically occurred after 1-2 weeks of culturing and sporophyte development took place after 3-4 weeks. These times are comparable to those reported for standard commercial nursery practices (37).

To evaluate the differential effects of bacteria on *Saccharina*, we calculated the log_2_fold-change for each well compared to the trial average for *Saccharina*-only controls for both the percent coverage of gametophytes and the number of sporophytes. We estimated gametophyte percent cover through microscopic assessment, documenting their approximate coverage percentage in the view. Sporophytes were also enumerated under microscopy as their distinct structural features [Fig. S4]. Log_2_fold-change cannot be calculated for values of 0 as was occasionally the case for sporophyte numbers, thus we added 1 to the number of sporophytes prior to comparison. We then used two-tailed t-tests followed by Benjamini-Hochberg correction for multiple comparisons to assess the differences between each bacterial isolate treatment and the *Saccharina*-only control group.

In trial 4 exclusively, 16S rRNA gene amplicon sequencing was employed to assess the bacterial community dynamics throughout the experiment. This allowed us to assess whether the bacterial cultures added on day 1 persisted over time. We sampled the microbiome of culture wells by swabbing the entire well destructively (culture media, well surfaces, and *Saccharina* in culture) at day 1, day 7, day 14, day 21, and day 28 for eight treatments, including the control.

### Indole Acetic acid screening

Indole acetic acid producing capabilities of 99 bacterial isolates were tested in batches of 10-20 using Salkowski reagent (ferric chloride + 35% perchloric acid). Isolates were plated on Difco Agar from glycerol stocks kept at -70℃ and allowed to grow for 2-3 days at room temperature. Following the appearance of bacterial colonies, isolates were transferred into Difco Marine Broth media and shaken at 130 rpm overnight. Bacterial cultures were then transferred into 0.15% (w/v) tryptophan media, covered in foil to block light and shaken at 130 rpm for 72 hours. After shaking, 1.5 mL of bacterial culture media was centrifuged for 5 minutes at 14,000 g, and 1 mL of the resulting supernatant was then mixed with 1 mL of Salkowski reagent. A standard curve of IAA concentrations was created for each batch of isolates tested by dissolving 0.05g of Indole-3-Acetic Acid (IAA) in 5 mL of 100% ethanol, then serially diluting in tryptophan media to final concentrations of 50, 40, 30, 20, 10, and 0 µg mL^-1^. One mL of these standard dilutions was mixed with 1 mL of Salkowski reagent and left in the dark alongside the isolate supernatant solutions for 30 minutes. After 30 minutes, 100 µL were transferred into a 96-well plate and spectrophotometric absorption measurements were taken at 536 nm. All isolates and standard curve dilutions were tested in duplicate. IAA producing ability was determined by comparing absorption measurements of isolates to the standard curve of IAA concentrations.

### Data Availability

Raw sequence files and metadata files have been deposited in the European Nucleotide Archive under the project accession PRJEB64485. All relevant raw data for bacterial isolates are within supplementary tables 1, 2, 3 and 4.

**Table 1.**
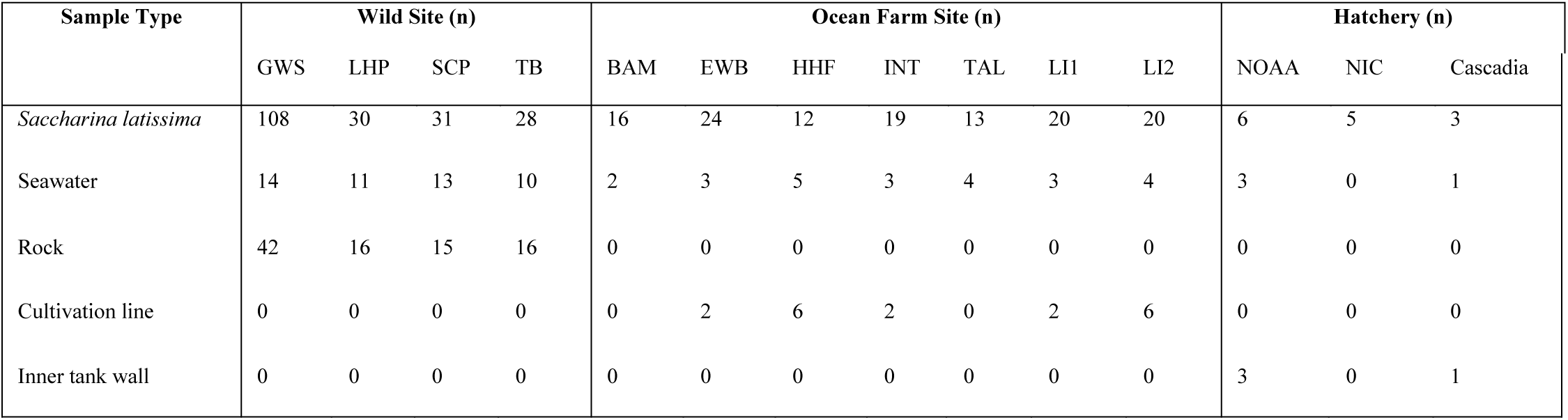
Overview of samples included in the datasets. . Bacterial samples included in our study sorted by sample type and location, including wild and cultivated sugar kelp (S. latissima) populations. The abbreviations represent sites (GWS = Girl in a Wet Suit, LHP = Lighthouse Park, SCP = Sandy Cove Park, and TB = Third Beach, BAM = Bamfield, EWB = East West Bay, HHF = Hood Head, INT = Interfor, TAL = Talbot Cove, LI = Loughboro Inlet, NOAA = WA nursery, NIC = BC nursery, Cascadia = BC nursery).

## ACKNOWLEDGMENTS

We respectfully acknowledge that this research was conducted on the traditional and unceded territory of the Coast Salish Peoples, including the Squamish (Skwxwú7mesh), Musqueam (xʷməθkʷəy̓əm), and Tsleil-Waututh (səl̓ilwətaɁɬ) Nations, upon whose land Stanley Park is situated.

J.P and L.W.P designed the study. J.P and S.S collected the wild kelp microbiome data at the GW site in Stanley Park, BC, Canada. S.S collected the wild kelp microbiome data from LHP, TB, and SCP. K.D and S.S also provided cultivated kelp microbiome data from the nursery and aquaculture, with support from J.C, a collaborator from Cascadia Seaweed. J.P conducted the molecular laboratory work for the partial wild kelp microbiome data. J.P. and E.K. carried out bioinformatics analyses. J.P isolated and identified all bacterial cultures used in the laboratory experiment. E.K conducted IAA experiments. With guidance from L.W.P, J.P wrote the manuscript and J.P and E.K created the figures. E.K, L.W.P, and J.P. revised the manuscript with input from all authors.

## References

1. Adams JM, Gallagher JA, Donnison IS. Fermentation study on *Saccharina latissima* for bioethanol production considering variable pre-treatments. J Appl Phycol. 2009;21(5):569.

2. Marinho GS, Holdt SL, Birkeland MJ, Angelidaki I. Commercial cultivation and bioremediation potential of sugar kelp, *Saccharina latissima*, in Danish waters. J Appl Phycol. 2015;27(5):1963–73.

3. Wargacki AJ, Leonard E, Win MN, Regitsky DD, Santos CNS, Kim PB, et al. An engineered microbial platform for direct biofuel production from brown macroalgae. Science. 2012;335(6066):308–13.

4. Bak UG, Nielsen CW, Marinho GS, Gregersen Ó, Jónsdóttir R, Holdt SL. The seasonal variation in nitrogen, amino acid, protein and nitrogen-to-protein conversion factors of commercially cultivated Faroese *Saccharina latissima*. Algal Res. 2019;42:101576.

5. Forbord S, Steinhovden KB, Solvang T, Handå A, Skjermo J. Effect of seeding methods and hatchery periods on sea cultivation of *Saccharina latissima* (Phaeophyceae): a Norwegian case study. J Appl Phycol. 2020;32(4):2201–12.

6. Peteiro C, Sánchez N, Martínez B. Mariculture of the Asian kelp *Undaria pinnatifida* and the native kelp *Saccharina latissima* along the Atlantic coast of Southern Europe: An overview. Algal Res. 2016 Apr 1;15:9–23.

7. Kim JK, Yarish C, Hwang EK, Park M, Kim Y. Seaweed aquaculture: cultivation technologies, challenges and its ecosystem services. ALGAE. 2017 Mar 15;32(1):1–13.

8. Augyte S, Yarish C, Redmond S, Kim JK. Cultivation of a morphologically distinct strain of the sugar kelp, *Saccharina latissima forma angustissima*, from coastal Maine, USA, with implications for ecosystem services. J Appl Phycol. 2017;29(4):1967–76.

9. Food FAO. Agriculture Organization of the United Nations. 2016. The State of World Fisheries and Aquaculture 2016. Contributing to food security and nutrition for all. Rome. 200 pp. 2018.

10. Coleman S, Gelais ATS, Fredriksson DW, Dewhurst T, Brady DC. Identifying Scaling Pathways and Research Priorities for Kelp Aquaculture Nurseries Using a Techno-Economic Modeling Approach. Front Mar Sci [Internet]. 2022 May 24 [cited 2024 Dec 20];9. Available from: https://www.frontiersin.org/journals/marine-science/articles/10.3389/fmars.2022.894461/full

11. Park J, Kim JK, Kong JA, Depuydt S, Brown MT, Han T. Implications of rising temperatures for gametophyte performance of two kelp species from Arctic waters. Bot Mar. 2017;60(1):39–48.

12. Steneck RS, Graham MH, Bourque BJ, Corbett D, Erlandson JM, Estes JA, et al. Kelp forest ecosystems: biodiversity, stability, resilience and future. Environ Conserv. 2002 Dec;29(4):436–59.

13. Ward GM, Faisan Jr JP, Cottier-Cook EJ, Gachon C, Hurtado AQ, Lim PE, et al. A review of reported seaweed diseases and pests in aquaculture in Asia. J World Aquac Soc. 2020;51(4):815–28.

14. Kim GH, Moon KH, Kim JY, Shim J, Klochkova TA. A revaluation of algal diseases in Korean *Pyropia* (Porphyra) sea farms and their economic impact. Algae. 2014;29(4):249–65.

15. Loureiro R, Gachon CMM, Rebours C. Seaweed cultivation: potential and challenges of crop domestication at an unprecedented pace. New Phytol. 2015;206(2):489–92.

16. Tsiresy G, Preux J, Lavitra T, Dubois P, Lepoint G, Eeckhaut I. Phenology of farmed seaweed *Kappaphycus alvarezii* infestation by the parasitic epiphyte *Polysiphonia sp*. in Madagascar. J Appl Phycol. 2016;28:2903–14.

17. Kouzuma A, Watanabe K. Exploring the potential of algae/bacteria interactions. Curr Opin Biotechnol. 2015;33:125–9.

18. Mancuso FP, D’Hondt S, Willems A, Airoldi L, De Clerck O. Diversity and Temporal Dynamics of the Epiphytic Bacterial Communities Associated with the Canopy-Forming Seaweed *Cystoseira compressa* (Esper) Gerloff and Nizamuddin. Front Microbiol. 2016;7:476.

19. Rosenberg E, Koren O, Reshef L, Efrony R, Zilber-Rosenberg I. The role of microorganisms in coral health, disease and evolution. Nat Rev Microbiol. 2007 May;5(5):355–62.

20. Sweet MJ, Bulling MT. On the Importance of the Microbiome and Pathobiome in Coral Health and Disease. Front Mar Sci. 2017;4.

21. Busby PE, Soman C, Wagner MR, Friesen ML, Kremer J, Bennett A, et al. Research priorities for harnessing plant microbiomes in sustainable agriculture. PLOS Biol. 2017 Mar 28;15(3):e2001793.

22. O’Callaghan M. Microbial inoculation of seed for improved crop performance: issues and opportunities. Appl Microbiol Biotechnol. 2016;100(13):5729–46.

23. Glick BR. Plant Growth-Promoting Bacteria: Mechanisms and Applications. Scientifica. 2012;2012:963401.

24. Bashan Y, Levanony H. Current status of *Azospirillum* inoculation technology: *Azospirillum* as a challenge for agriculture. Can J Microbiol. 1990 Sep;36(9):591–608.

25. Patten CL, Glick BR. Role of *Pseudomonas putida* Indoleacetic Acid in Development of the Host Plant Root System. Appl Environ Microbiol. 2002 Aug;68(8):3795.

26. Amin SA, Hmelo LR, van Tol HM, Durham BP, Carlson LT, Heal KR, et al. Interaction and signalling between a cosmopolitan phytoplankton and associated bacteria. Nature. 2015 Jun;522(7554):98–101.

27. Park J, Park BS, Wang P, Patidar SK, Kim JH, Kim SH, et al. Phycospheric native bacteria *Pelagibaca bermudensis* and *Stappia sp*. ameliorate biomass productivity of T*etraselmis striata* (KCTC1432BP) in co-cultivation system through mutualistic interaction. Front Plant Sci. 2017;8:289.

28. Wang H, Elyamine AM, Liu Y, Liu W, Chen Q, Xu Y, et al. *Hyunsoonleella sp.* HU1-3 Increased the Biomass of *Ulva fasciata*. Front Microbiol. 2022;12:788709.

29. Dittami SM, Duboscq-Bidot L, Perennou M, Gobet A, Corre E, Boyen C, et al. Host– microbe interactions as a driver of acclimation to salinity gradients in brown algal cultures. ISME J. 2016;10(1):51–63.

30. Ma M, Zhuang Y, Chang L, Xiao L, Lin Q, Qiu Q, et al. Naturally occurring beneficial bacteria Vibrio alginolyticus X-2 protects seaweed from bleaching disease. Mbio. 2023;e00065–23.

31. Singh RP, Bijo AJ, Baghel RS, Reddy CRK, Jha B. Role of bacterial isolates in enhancing the bud induction in the industrially important red alga *Gracilaria dura*. FEMS Microbiol Ecol. 2011;76(2):381–92.

32. Spoerner M, Wichard T, Bachhuber T, Stratmann J, Oertel W. Growth and Thallus Morphogenesis of *Ulva mutabilis* (Chlorophyta) Depends on A Combination of Two Bacterial Species Excreting Regulatory Factors. J Phycol. 2012;48(6):1433–47.

33. Tapia JE, González B, Goulitquer S, Potin P, Correa JA. Microbiota Influences Morphology and Reproduction of the Brown Alga Ectocarpus sp. Front Microbiol [Internet]. 2016 [cited 2023 May 5];7. Available from: https://www.frontiersin.org/articles/10.3389/fmicb.2016.00197

34. Wichard T. Exploring bacteria-induced growth and morphogenesis in the green macroalga order Ulvales (Chlorophyta). Front Plant Sci. 2015;6:86.

35. Li J, Majzoub ME, Marzinelli EM, Dai Z, Thomas T, Egan S. Bacterial controlled mitigation of dysbiosis in a seaweed disease. ISME J. 2022;16(2):378–87.

36. Redmond S, Green L, Yarish C, Kim J, Neefus C. New England Seaweed Culture Handbook-Nursery Systems [Internet]. Connecticut Sea Grant; 2014. Available from: http://seagrant.uconn.edu/publications/aquaculture/handbook.pdf

37. Forbord S, Steinhovden KB, Rød KK, Handå A, Skjermo J. Cultivation protocol for *Saccharina latissima*. Protoc Macroalgae Res. 2018;2:4.1.

38. Flavin K, Flavin N, Flahive B. Kelp farming manual: a guide to the processes, techniques, and equipment for farming kelp in New England waters. Ocean Approv LLC Saco. 2013;

39. Rolin C, Inkster R, Laing J, Hedges J, McEvoy L. Seaweed cultivation manual. Shetl Seaweed Grow Proj 2014. 2016;16.

40. Yarish C, Kim JK, Lindell S, Kite-Powell H. Developing an environmentally and economically sustainable sugar kelp aquaculture industry in southern New England: from seed to market. 2017;

41. Lian J, Wijffels RH, Smidt H, Sipkema D. The effect of the algal microbiome on industrial production of microalgae. Microb Biotechnol. 2018 Jul 5;11(5):806–18.

42. Marshall K, Joint I, Callow ME, Callow JA. Effect of Marine Bacterial Isolates on the Growth and Morphology of Axenic Plantlets of the Green Alga *Ulva linza*. Microb Ecol. 2006 Aug 1;52(2):302–10.

43. Fries L. Axenic tissue cultures from the sporophytes of *Laminaria digitata* and *Laminaria hyperborea* (Phaeophyta). J Phycol. 1980;16(3):475–7.

44. Morris MM, Haggerty JM, Papudeshi BN, Vega AA, Edwards MS, Dinsdale EA. Nearshore Pelagic Microbial Community Abundance Affects Recruitment Success of Giant Kelp, Macrocystis pyrifera. Front Microbiol [Internet]. 2016 Nov 14 [cited 2024 Sep 4];7. Available from: https://www.frontiersin.org/journals/microbiology/articles/10.3389/fmicb.2016.01800/full

45. Osborne MG, Simons AL, Molano G, Tolentino B, Singh A, Arismendi GJM, et al. Investigating the relationship between microbial network features of giant kelp “seedbank” cultures and subsequent farm performance. PLOS ONE. 2024 Mar 27;19(3):e0295740.

46. Burgunter-Delamare B, Rousvoal S, Legeay E, Tanguy G, Fredriksen S, Boyen C, et al. The *Saccharina latissima* microbiome: Effects of region, season, and physiology. Front Microbiol. 2023;13:1050939.

47. Phelps CM, McMahon K, Bissett A, Bernasconi R, Steinberg PD, Thomas T, et al. The surface bacterial community of an Australian kelp shows cross-continental variation and relative stability within regions. FEMS Microbiol Ecol. 2021;97(7):fiab089.

48. Schenk S, Wardrop CG, Parfrey LW. Low salinity destabilizes the bacterial community of sugar kelp, Saccharina latissima (Phaeophyceae) [Internet]. bioRxiv; 2024 [cited 2025 Feb 7]. p. 2023.12.07.570704. Available from: https://www.biorxiv.org/content/10.1101/2023.12.07.570704v3

49. King NG, Moore PJ, Thorpe JM, Smale DA. Consistency and Variation in the Kelp Microbiota: Patterns of Bacterial Community Structure Across Spatial Scales. Microb Ecol. 2023 May 1;85(4):1265–75.

50. Marzinelli EM, Campbell AH, Zozaya Valdes E, Vergés A, Nielsen S, Wernberg T, et al. Continental-scale variation in seaweed host-associated bacterial communities is a function of host condition, not geography. Environ Microbiol. 2015;17(10):4078–88.

51. Björk JR, O’Hara RB, Ribes M, Coma R, Montoya JM. The dynamic core microbiome: Structure, dynamics and stability. bioRxiv. 2018;137885.

52. Neu AT, Allen EE, Roy K. Defining and quantifying the core microbiome: Challenges and prospects. Proc Natl Acad Sci. 2021;118(51):e2104429118.

53. Shade A, Handelsman J. Beyond the Venn diagram: the hunt for a core microbiome. Environ Microbiol. 2012;14(1):4–12.

54. Hernandez-Agreda A, Leggat W, Bongaerts P, Herrera C, Ainsworth TD. Rethinking the Coral Microbiome: Simplicity Exists within a Diverse Microbial Biosphere. mBio. 2018 Oct 9;9(5):10.1128/mbio.00812-18.

55. Risely A. Applying the core microbiome to understand host–microbe systems. J Anim Ecol. 2020 Jul 1;89(7):1549–58.

56. Shade A, Stopnisek N. Abundance-occupancy distributions to prioritize plant core microbiome membership. Curr Opin Microbiol. 2019 Jun;49:50–8.

57. Davis KM, Zeinert L, Byrne A, Davis J, Roemer C, Wright M, et al. Successional dynamics of the cultivated kelp microbiome. J Phycol. 2023;

58. Gang S, Sharma S, Saraf M, Buck M, Schumacher J. Analysis of Indole-3-acetic Acid (IAA) Production in *Klebsiellaby* LC-MS/MS and the Salkowski Method. Bio-Protoc. 2019 May 5;9(9):e3230.

59. Ghaderiardakani F, Coates JC, Wichard T. Bacteria-induced morphogenesis of *Ulva intestinalis* and *Ulva mutabilis* (Chlorophyta): a contribution to the lottery theory. FEMS Microbiol Ecol. 2017 Aug 1;93(8):fix094.

60. Kessler RW, Weiss A, Kuegler S, Hermes C, Wichard T. Macroalgal–bacterial interactions: Role of dimethylsulfoniopropionate in microbial gardening by *Ulva* (Chlorophyta). Mol Ecol. 2018;27(8):1808–19.

61. Louca S, Polz MF, Mazel F, Albright MBN, Huber JA, O’Connor MI, et al. Function and functional redundancy in microbial systems. Nat Ecol Evol. 2018 Jun;2(6):936–43.

62. Haney CH, Wiesmann CL, Shapiro LR, Melnyk RA, O’Sullivan LR, Khorasani S, et al. Rhizosphere-associated Pseudomonas induce systemic resistance to herbivores at the cost of susceptibility to bacterial pathogens. Mol Ecol. 2018;27(8):1833–47.

63. Lin JD, Lemay MA, Parfrey LW. Diverse Bacteria Utilize Alginate Within the Microbiome of the Giant Kelp Macrocystis pyrifera. Front Microbiol [Internet]. 2018 Aug 20 [cited 2024 Dec 20];9. Available from: https://www.frontiersin.org/journals/microbiology/articles/10.3389/fmicb.2018.01914/full

64. Martiny JBH, Jones SE, Lennon JT, Martiny AC. Microbiomes in light of traits: A phylogenetic perspective. Science. 2015 Nov 6;350(6261):aac9323.

65. Hehemann JH, Arevalo P, Datta MS, Yu X, Corzett CH, Henschel A, et al. Adaptive radiation by waves of gene transfer leads to fine-scale resource partitioning in marine microbes. Nat Commun. 2016 Sep 22;7(1):12860.

66. Gu K, Chen CY, Selvaraj P, Pavagadhi S, Yeap YT, Swarup S, et al. *Penicillium citrinum* Provides Transkingdom Growth Benefits in Choy Sum (*Brassica rapa var. parachinensis*). J Fungi. 2023 Apr;9(4):420.

67. Spaepen S, Vanderleyden J. Auxin and Plant-Microbe Interactions. Cold Spring Harb Perspect Biol. 2011 Apr;3(4):a001438.

68. Zhang S, van Duijn B. Cellular Auxin Transport in Algae. Plants. 2014 Jan 27;3(1):58–69.

69. Chen X, Tang Y, Sun X, Zhang X, Xu N. Comparative transcriptome analysis reveals the promoting effects of IAA on biomass production and branching of *Gracilariopsis lemaneiformis*. Aquaculture. 2022 Feb 15;548:737678.

70. Kai T, Nimura K, Yasui H, Mizuta H. Regulation of Sorus Formation by Auxin in Laminariales Sporophyte. J Appl Phycol. 2006 Feb 1;18(1):95–101.

71. Lee JH, Wood TK, Lee J. Roles of Indole as an Interspecies and Interkingdom Signaling Molecule. Trends Microbiol. 2015 Nov 1;23(11):707–18.

72. Zarkan A, Liu J, Matuszewska M, Gaimster H, Summers DK. Local and Universal Action: The Paradoxes of Indole Signalling in Bacteria. Trends Microbiol. 2020 Jul 1;28(7):566–77.

73. Albright MBN, Louca S, Winkler DE, Feeser KL, Haig SJ, Whiteson KL, et al. Solutions in microbiome engineering: prioritizing barriers to organism establishment. ISME J. 2022 Feb 1;16(2):331–8.

74. Poppeliers SW, Sánchez-Gil JJ, de Jonge R. Microbes to support plant health: understanding bioinoculant success in complex conditions. Curr Opin Microbiol. 2023 Jun 1;73:102286.

75. Caporaso JG, Lauber CL, Walters WA, Berg-Lyons D, Huntley J, Fierer N, et al. Ultra-high-throughput microbial community analysis on the Illumina HiSeq and MiSeq platforms. ISME J. 2012 Aug;6(8):1621–4.

76. Fadrosh DW, Ma B, Gajer P, Sengamalay N, Ott S, Brotman RM, et al. An improved dual-indexing approach for multiplexed 16S rRNA gene sequencing on the Illumina MiSeq platform. Microbiome. 2014;2(1):1–7.

77. Quast C, Pruesse E, Yilmaz P, Gerken J, Schweer T, Yarza P, et al. The SILVA ribosomal RNA gene database project: improved data processing and web-based tools. Nucleic Acids Res. 2012;41(D1):D590–6.

78. Callahan BJ, McMurdie PJ, Rosen MJ, Han AW, Johnson AJA, Holmes SP. DADA2: high-resolution sample inference from Illumina amplicon data. Nat Methods. 2016;13(7):581.

79. Rosen MJ, Callahan BJ, Fisher DS, Holmes SP. Denoising PCR-amplified metagenome data. BMC Bioinformatics. 2012 Oct 31;13(1):283.

80. McMurdie PJ, Holmes S. phyloseq: an R package for reproducible interactive analysis and graphics of microbiome census data. PloS One. 2013;8(4):e61217.

81. Bray JR, Curtis JT. An ordination of the upland forest communities of southern Wisconsin. Ecol Monogr. 1957;27(4):325–49.

82. Oksanen J, Blanchet FG, Friendly M, Kindt R, Legendre P, McGlinn D, et al. vegan: Community Ecology Package. R package version 2.4–6. 2018. 2019.

83. De Caceres M, Jansen F, De Caceres MM. Package ‘indicspecies.’ indicators. 2016;8:1.

84. Cáceres MD, Legendre P. Associations between species and groups of sites: indices and statistical inference. Ecology. 2009;90(12):3566–74.

85. Legendre P. Indicator species: computation. 2013;

86. Park J, Davis K, Lajoie G, Parfrey LW. Alternative approaches to identify core bacteria in Fucus distichus microbiome and assess their distribution and host-specificity. Environ Microbiome. 2022;17(1):1–15.

87. Wickham H. ggplot2: elegant graphics for data analysis (use R!). springer New York; 2009.

88. Lane DJ. 16S/23S rRNA sequencing. Nucleic acid techniques in bacterial systematics (Stackebrandt E & Goodfellow M, eds). Wiley, New York; 1991.

89. Madden T. The BLAST sequence analysis tool. NCBI Handb. 2003;

90. Bolyen E, Rideout JR, Dillon MR, Bokulich NA, Abnet CC, Al-Ghalith GA, et al. Reproducible, interactive, scalable and extensible microbiome data science using QIIME 2. Nat Biotechnol. 2019;37(8):852–7.

91. Letunic I, Bork P. Interactive Tree Of Life (iTOL): an online tool for phylogenetic tree display and annotation. Bioinformatics. 2007;23(1):127–8.

92. Sutton S. Measurement of microbial cells by optical density. J Valid Technol. 2011;17(1):46– 9.

